# All^2^: A tool for selecting mosaic mutations from comprehensive multi-cell comparisons

**DOI:** 10.1101/2021.09.29.462281

**Authors:** Vivekananda Sarangi, Yeongjun Jang, Milovan Suvakov, Taejeong Bae, Liana Fasching, Shobana Sekar, Livia Tomasini, Jessica Mariani, Flora M. Vaccarino, Alexej Abyzov

## Abstract

Accurate discovery of somatic mutations in a cell is a challenge that partially lays in immaturity of dedicated analytical approaches. Approaches comparing cell’s genome to a control bulk sample miss common mutations, while approaches to find such mutations from bulk suffer from low sensitivity. We developed a tool, All^2^, which enables accurate filtering of mutations in a cell from exhaustive comparison of cells’ genomes to each other without data for bulk(s). Based on all pair-wise comparisons, every variant call (point mutation, indel, and structural variant) is classified as either a germline variant, mosaic mutation, or false positive. As All^2^ allows for considering dropped-out regions, it is applicable to whole genome and exome analysis of cloned and amplified cells. By applying the approach to a variety of available data, we showed that its application reduces false positives, enables sensitive discovery of high frequency mutations, and is indispensable for conducting high resolution cell lineage tracing. All^2^ is freely available at https://github.com/abyzovlab/All2.

## Introduction

With advances in sequencing technologies, analysis of the genome of a single cell is gaining traction owing to its multiple applications, including studying mutations in normal and cancer single cells, identification of driver mutations in cancer (1, 2), tracing cell lineages in human development (3, 4) and identification of sub-clones in cancer (1, 2), where mosaic mutations are used as barcodes to study cell lineages. For detection of mosaic mutations in single cells, the most frequently used approach is to compare single cell genomes to that of a matched reference bulk. While this approach works well to find private mutations in a cell, it misses mutations that are present at higher frequency, and consequently present in multiple cells in the reference bulk. It also requires one to have bulk data which might not be always available. Here we present a tool called All^2^ (pronounced ‘all square’) which detects mosaic mutations without the need for a reference bulk by relying on comprehensive cell-to-cell comparisons. By consolidating information from all comparisons, every call is categorized as either a germline variant, mosaic mutation or noise/false positive.

## Results

### Concept

All^2^ is an easy-to-use tool which extends and implements an algorithm initially proposed in Bae et al. 2018 (5). All^2^ takes mutation calls from all pair-wise comparisons of *N* cells in the study and, for every non-redundant call, creates a *NxN* pairwise binary matrix corresponding to comparing different pairs of cells, where 1 corresponds to a call and 0 to no call. Patterns of values in the matrix are used to determine whether a call is a mosaic mutation, germline variant or false positive (**Fig. 1A-D**). In theory, the presence of these patterns should be sufficient to make the determination, however, real data has noise, smearing the patterns (**Fig. S1**).

**Figure 1.**
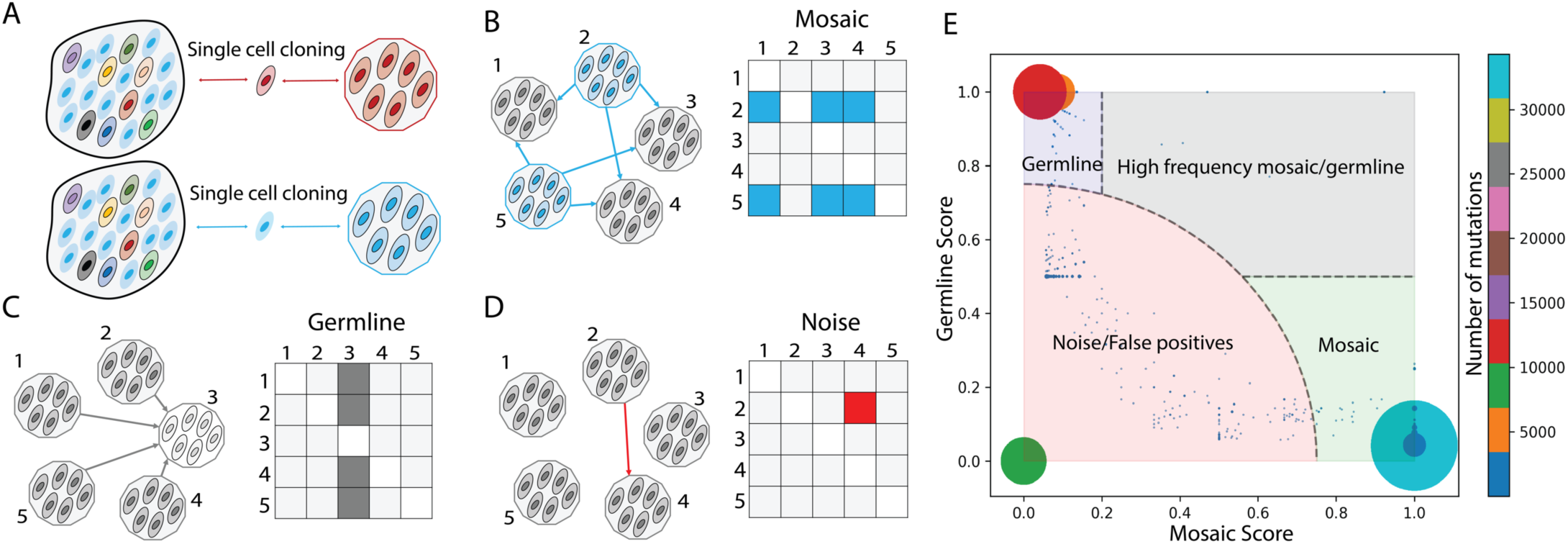
Conceptual overview of All^2^ approach and scoring. **A)** A tissue/sample is made up of different cells (ovals) carrying various mosaic mutations (reflected by different colors). Post single cell clonal expansion, rare mosaic mutations (in red) can be easily detected by comparing the clone to the bulk tissue. However, frequent mutations (in blue) will be missed by this approach. **B-D)** Each mutation in clone-to-clone (which is cell-to-cell) comparison can be represented by a NxN matrix of pairwise clone comparisons, where each box represents the call between a clone in the row versus a clone in the column. **B)** In case of a true mosaic mutation, the calls are arranged as rows in the matrix. The pattern in the matrix shows that the mutation is called in clone 2 and clone 5 when comparing them to other clones. **C)** In case of a germline variant, the calls are arranged in a column in the matrix. The displayed pattern suggests that the mutation is present in all clones except clone 3. **D)** The pattern has a sporadic distribution of calls in the pairwise matrix and does not suggest either mosaic mutations or germline variants. Such call is deemed as a false positive or noise. **E)** Distribution of mosaic and germline scores for calls (the size of the dot/circle corresponds to the number of calls with the same scores). The plot can be divided into four areas: mosaic mutations (where the mutations have high mosaic scores and low germline scores), germline variants (where the mutations have high germline and low mosiac scores), high frequency mosaic mutations (where calls have high both mosaic and germline scores) and, lastly, noise or false positive calls.

For effective categorization, we developed a scoring system which reflects how likely it is for a call to be a mosaic mutation or a germline variant. The tool calculates two scores: a germline score and a mosaic score, each within a range between 0 and 1. A real mosaic variant could only be discovered when comparing a cell carrying the variant and a cell not carrying one. The number of times a call for a variant shows up in the matrix is determined by

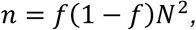

where *n* is the number of times a variant is seen in all comparison, *f* is fraction of cells carrying the variant, *N* is the total number of cells. By solving the above quadratic equation, we get two solutions:

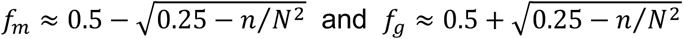

Since the mosaic mutations are typically present in a small fraction of cells in the bulk, and germline variants are present in (almost) all the cells, we conditionally call *f*_*m*_ as frequency of a mosaic mutation and *f*_*g*_ as frequency of a germline variant. Note, a germline variant can be lost or undetected in some cells, and that is why its cell frequency in a bulk may be below 1.

Since *f*_*m*_ = 1 –*f*_*g*_, we can just use one, such as *f = f*_*m*_, where *f* is the fraction of cells with mosaic mutation or the fraction of cells without germline variant. Now, we can calculate the number of cells *N′* carrying the mosaic mutation or the number of cells not carrying germline variants as *N′ ≈ fN*. In case of a true mosaic mutation, the corresponding calls are arranged in rows in the matrix (**Fig. 1B**), and would sum up to

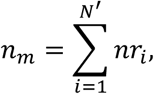

where *nr*_*i*_ is the number of calls for the variant in a row corresponding to the *i*^*th*^ cell. From the data, the best estimate of *n*_*m*_ is the maximum from all possible subset of *N’* cells from *N*. Similarly, for a germline variant, corresponding calls are arranged in columns (**Fig. 1C**), and would sum up to

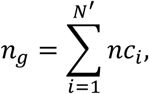

where *nc*_*i*_ is the number of calls for the variant in a column corresponding to *i*^*th*^ cell. And best estimate of *n*_*g*_ is the maximum from all possible subset of *N’* cells from *N*. The mosaic and germline scores are then defined as

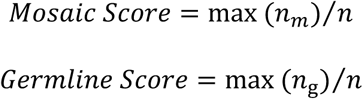

A call having a high mosaic score and low germline score is defined as a mosaic mutation. Similarly, a call with high germline score and low mosaic score is defined as a germline score. When a call has both high germline and mosaic scores, we define it as a high frequency mosaic mutation. Such mutations are likely present at a higher fraction of cells in a tissue, yet at a lower fraction than germline variant. For example, such mutations could occur during early development and be present in a high fraction of cells across tissues in the human body (6). The distribution of mutations (as dots) on a plane with axes corresponding to the two scores can be used to divide the calls into mosaic mutations, germline variants, noise or false positive (low mosaic and low germline score) and high frequency mosaic mutations (high mosaic and high germline score) (**Fig. 1E)**.

#### Implementation and usage

Genomes of all pairs of cells need to be compared prior to using All^2^. Variant calls can be made by a caller of choice (see **Methods**). All^2^ is written in python and has three commands: ‘score’, ‘call’, and ‘matrix’. The first command takes a manifest file with names of single cells, along with the VCF file containing calls (SNVs and INDELs) as rows. Case and control fields in the manifest file are used to define the directions of pairwise comparisons, where the case is compared to control. A user can optionally provide a BED file with the inclusion list of genomic regions where to apply the filtering (see **Applications**). The output of this command is a file with mosaic and germline scores for each of the calls as well as density scatter plot of the scores showing distribution of calls based on their scores (**Fig. S2A**). The second command relies on the output of the ‘score’ command and annotates the calls as mosaic mutation, germline variant, noise, or high frequency mosaic mutations based on default or user specified score cut-offs. This command also annotates the density scatter plot (**Fig. 1E**), provides a file with annotated calls for each cell, per category plots of call counts, per sample plots of call counts, VAF (variant allele frequency) plot, and mutation spectrum plots (**Fig. S2B-D**). The third command plots a matrix of pairwise comparison for one or multiple calls. The plots also display calculated scores along with VAF for the call(s) in each cell (**Fig. S3**). Analogous to SNVs and indels, All^2^ is capable of filtering structural variant (SV) calls using commands ‘score_sv’, ‘call_sv’ and ‘matrix_sv’. Two SV calls are considered the same if they have at least 50% reciprocal overlap. For this purpose, the tool supports VCF file as input, e.g., VCFs generated by the SV caller MANTA (7).

One implicit underlying assumption of the approach is that in each compared cell, the genome is covered/sequenced uniformly. This is true in case of the single cell cloning approach, however, single cell genome amplification may result in non-uniform coverage which, at the extreme, manifests in allelic dropouts (8). To handle this, we have implemented a dedicated allele dropout analysis (ADA) mode, which considers allele dropout regions when calculating the scores, thereby reducing the noise (see **Application**). The ADA mode can also be used for running All^2^ on exome data where the exome capture region can be specified per cell in the manifest file. More details of the command and description of the results can be found on the dedicated GitHub page https://github.com/abyzovlab/All2.

#### Application to reconstructing cell lineage tree

To demonstrate the uniqueness of All^2^ approach, we applied it to reconstruct post-zygotic cell divisions in a living individual. Analysis of developmental cell lineages is one of the central questions in developmental biology, resolving which can shed light on the etiology of developmental diseases. Unlike model organisms, lineage tracing in humans can only be done retrospectively using naturally occurring somatic variants that serve as permanent marks of the lineages. Mutations that occur during early development are present in a high fraction of cells across tissues in the human body, and their discovery is challenging for existing methods.

In the analyzed individual, we compared mosaic variant discovery using three approaches: 1) by analysis of bulk blood and saliva; 2) by pairwise comparison of 25 clonal iPSC lines (representing 25 fibroblast single cells) with the bulk blood; and 3) by comparing the clonal lines followed by application of All^2^. To reconstruct the lineage tree, we selected mosaic variants shared by clones or by multiple bulk tissues (6) (**Fig. 2**). Analysis of bulks alone allowed discovering only high frequency mutations but not all. For example, mutations *a, b*, and *c* defining branches of the first zygotic cleavage (**Fig. 2B**) could not be discovered because of resembling germline variants by frequency of occurrence in the bulks (i.e., in 80% to 90% of cells). Pairwise comparisons between clones and bulk tissues are powered to find mutations present in the analyzed cell and at low frequency (typically <1% VAF) in bulks but misses high frequency mutations. Remarkably, the All^2^ approach was able to call both high and low frequency mutations resulting in the most complete lineage tree – a tree that cannot be reconstructed even if we combine comparisons of clones relative to bulk tissues and analysis of bulks. The advantage of calling mosaic mutations in bulk is that it allows discovering mutations with intermediate VAF (between 1% to 10%), which were not sampled by the 25 analyzed clones and consequently, not discovered by All^2^. In fact, most mutations discovered from bulks were not sampled by the clones. Increasing the number of analyzed clones will likely increase the overlap in discovered mutations between those two approaches but would also increase experimental cost. Thus, this observation suggests complementarity in analyzing clones/single cells and bulks for lineage reconstruction.

**Figure 2.**
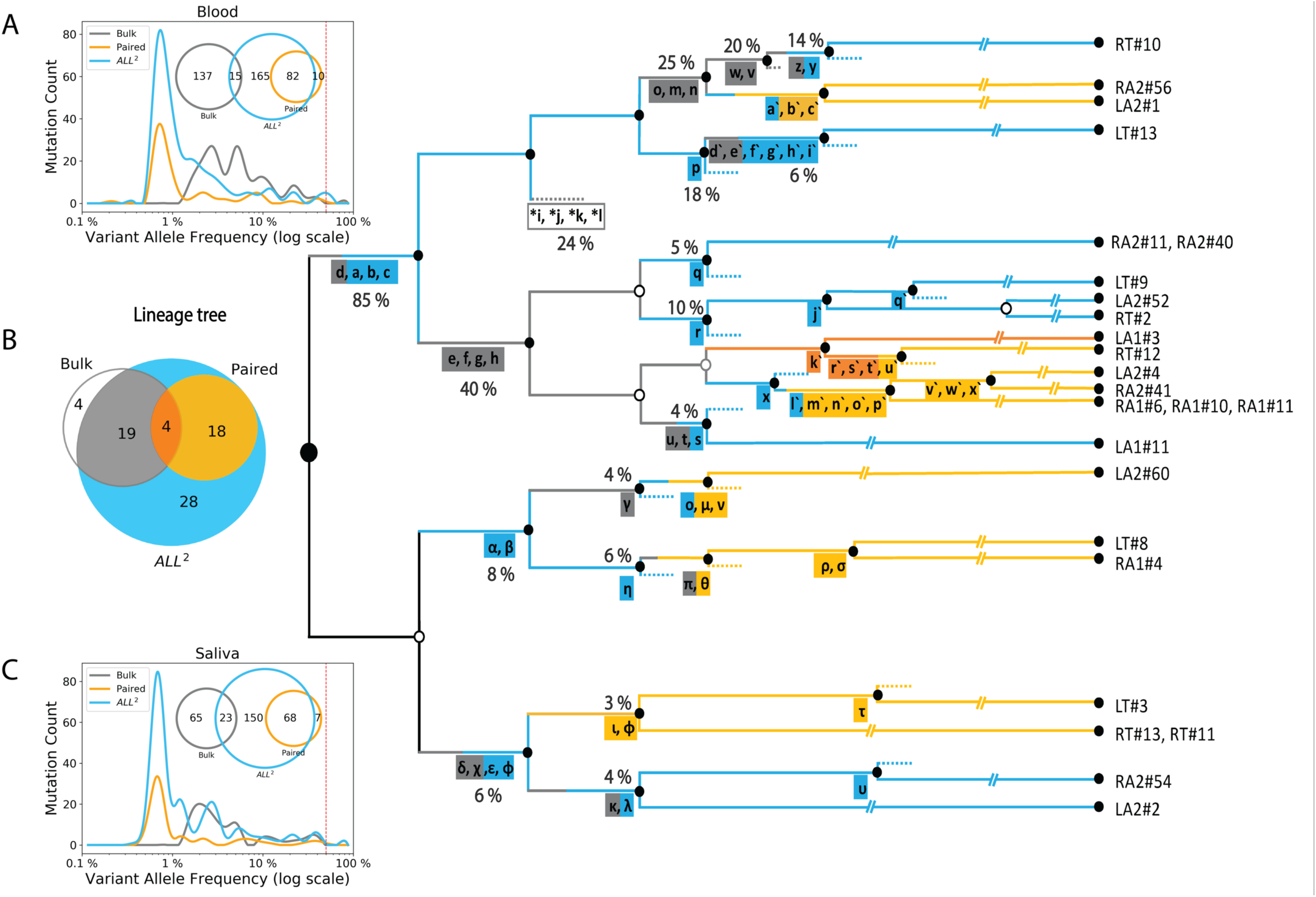
Calls from All^2^ enable reconstruction of high-resolution lineage tree. **A, C)** Application of All^2^ to iPSC clones discovers more variants (cyan) than analysis of deeply sequenced bulk tissues (gray) or pairwise comparison of clonal lines and the bulk (orange). The approach also calls variants across entire VAF spectrum. Analysis of bulk may discover variants with intermediate VAF (1%-10%) which are not sampled in clones. For the displayed comparison, variants with at least two supporting reads in the bulks are considered for each discovery approach. **B)** Lineage tree reconstructed from the analysis of 25 clones from an adult individual. Variants discovered from either bulk (gray) or pairwise (orange) comparisons provide limited information as compared to All^2^, which is the most comprehensive. Multiple additional branches in the lineage tree can be traced when using additional variants (cyan) discovered by applying only the All^2^ approach, which is also reflected in the Venn diagram. SNVs found only in the bulk tissues are marked with asterisks and define putative branch not sampled by clones. The percentage values next to branches denote the average fraction of the bulk cells carrying the mutations. Clone names are shown on the right.

#### Allele dropout mode for whole genome amplified single cells

Using clones as gold standard, we applied All^2^ in Allele Dropout Analysis (ADA) mode (see Methods) to MDA amplified single cells, to demonstrate the effectiveness of this mode to filter out spurious calls originating from biases in the amplification process. MDA uses φ29 polymerase under isothermal condition, which results in an exponential DNA amplification. The exponential amplification leads to uneven coverage and over representation of one allele over the other (allele imbalance). In extreme, a locus can have only one allele amplified and germline variants on the unamplified locus will be lost. ADA mode is designed to address this issue. In ADA mode, All^2^ takes a list of genomic regions (in bed format) where no allele dropout is observed (see **Methods**). Using this, for each call, All^2^ excludes from the score calculation those cells where a call is not made, and the surrounding region has allele dropout. This exclusion may change the number of considered cells and pairwise comparisons, which eventually affects the mosaic and germline scores.

We called mosaic mutations in 11 clones (representing 11 brain progenitor cells) derived from a human fetal brain (specimen 316) (5), as well as in an MDA amplified single cell taken from one of the clones. Just by adding the single MDA-amplified cell into the analysis, more than doubled the mutation counts (**Fig. 3A, B**). Next, we applied a single cell specialized caller named SCOUT(9). We observed that even though this partially reduced the effect of MDA amplification artifacts (**Fig. 3C**), it still resulted in a large number of mosaic and high frequency mosaic mutations, which was further mitigated by the ADA mode (**Fig. 3D**). Additionally, there is a reduction in the germline variants after applying ADA. These mutations, falsely called as homozygous reference due to allele dropout in the single cell, are effectively filtered by the ADA mode. Mutation counts per clone (**Fig. 3D**) were also similar to those found when analyzing only clones (**Fig. 3A**). This comparison yields evidence that even though the number of mutations called in the single cell is high, by applying ADA mode, we were able to reduce the number of potential false calls introduced by single cell amplification by half, without compromising the mutation calls from clones not affected by allele dropout.

**Figure 3.**
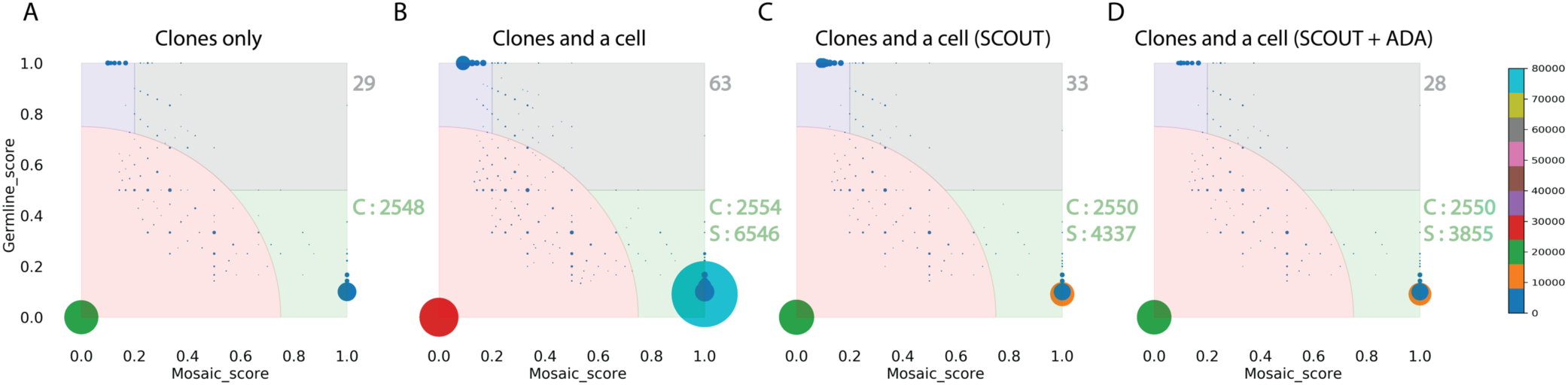
All^2^ in ADA mode reduces false positive calls from allele dropout in MDA. **A)** Score distribution when applying All^2^ to 11 clones derived from single brain progenitor cells. There are 29 calls for high frequency (gray area) and 2548 calls for low frequency (green area) mosaic mutations. The ‘C’ points to mosaic calls in the clones). **B)** Adding one MDA amplified cell to the analysis results in double the number of calls for high frequency mosaic mutations. Noise also increases. The ‘S’ points to the calls coming from the single cell. **C)** Application of a specialized single cell caller SCOUT on the single cell partially mitigates issues with calling, i.e., reduces the noise and the number of mosaic calls. **D)** Applying the ADA mode results in almost the same set of high frequency mosaic mutations. The mode also reduced calls for mosaic mutations in single cell without affecting calls in the clones.

#### Runtime

Runtime depends on the number of cells in the study and the variant caller used (since some variant callers will output higher number of calls than others). For the first example with 25 iPSC lines (**Fig. 2B**), application of All^2^ using 8 GB memory on a 2.4 GHz dual-core processor took less than 15 minutes for the ‘score’ module and less than 10 minutes for the ‘call’ module to compute mutation annotation and plot the mutation count and VAF plots. For the second example with 11 clones and one single cell (**Fig. 3**), application of All^2^ in ADA mode took less than 90 minutes to complete the ‘score’ module and less than 20 minutes for the ‘call’ module. In this case, the runtime is longer because of the longer list of variant candidates from the single cell.

## Conclusion

We have developed and implemented All^2^, which can discover mosaic SNVs, indels, and SVs from exhaustive cell-to-cell comparison of WGS data from single cells or clones. Our method is superior to using deep sequencing of bulk tissues and/or paired comparison of single cells versus bulk for detection of both low and high frequency mosaic mutations. A limitation relative to bulk method is that the mutations that are not sampled by the analyzed single cells cannot be discovered. This can be addressed by increasing the number of analyzed single cell. We have also applied All^2^ for comprehensive reconstruction of a developmental lineage tree, showing that All^2^ allows a vastly more comprehensive lineage discovery. Furthermore, the method is general and can be applied to any problem of lineage tracing that relies on the analysis of multiple cells, such as tracing cancer evolution.

We further demonstrate that All^2^ facilitates removal of false positive calls (in ADA mode) from amplified single cells. Additionally, since ADA mode takes a bed file with inclusive regions as input, All^2^ can be applied to the analyses of exome sequencing where a user can provide a file with target regions. The same mode can also be applied to exclude copy number altered regions when analyzing cancer cells. All^2^ provides visualizations such as allele frequency distribution, mutation spectrum, mutation counts and score distribution plots to help guide the user to better understand their data as well as change parameter setting for calling mosaic mutations. The tool is open source and is freely available on GitHub: https://github.com/abyzovlab/All2.

## Methods

### iPSC line generation

The iPSC lines were derived from fibroblasts using the Epi5 Episomal iPSC Reprogramming Kit (Invitrogen catalog A15960) delivering the five reprogramming factors Oct4, Sox2, Klf4, L-Myc, and Lin28. The iPSC lines were propagated using mTeSR1 media (Stem Cell Technologies) on 1X Matrigel-coated dishes (Matrigel®). Genomic DNA was extracted at passage six, using QIamp DNA Minikit (Qiagen) following the manufacturer instructions.

### Saliva collection and DNA extraction

Saliva DNA was collected and purified using the Oragene-Discover kit (DNA Genotek) following the manufacturer instructions. Saliva DNA was extracted using DNeasy Blood and Tissue kit (Qiagen) with the following modifications: 5 ml AL-buffer and 200 μl Proteinase K were added to saliva and incubated at 56°C for 30 minutes. RNA was digested using 20μl RNAse A (Qiagen) for 5 minutes and DNA was extracted using 4 extraction columns in parallel to optimize the yield.

### Blood collection and DNA extraction

10-15 ml of blood was collected using BD Vacutainer ACD tubes. DNA was extracted using the Gentra Puregene Blood Kit (Qiagen) following standard manufacturer protocols.

### Whole genome sequencing (WGS)

DNA extracted from iPSC lines were sequenced at 30X, while DNA extracted from saliva and blood was sequenced at 200X. All sequencing was conducted at BGI using with 2×100 bp paired reads. The sequencing library preparation was PCR-free.

### Fetal brain tissue and MDA

Collection of fetal brain tissues for subject 316, derivation of clonal neurosphere lines and sequencing has been previously described (5). Single cells from a clonal neurosphere line were manually picked using a micropipette under an inverted microscope. Whole genome amplification was performed by multiple displacement amplification (MDA) using the REPLI-g Single Cell Kit (QIAGEN) following the manufacturer recommendations. Genomic DNA was extracted using the DNeasy Blood & Tissue Kit (QIAGEN). Multiplex PCR for four arbitrary loci from different chromosomes was used to exclude single cells if less than four loci were amplified (10). Five out of eight single cells (62.5%) passed the 4-locus multiplex PCR quality control and were selected for sequencing. Illumina Truseq DNA PCR-free libraries were prepared for the five cells and sequenced on a HiSeq X (2×150 bp) at 30X coverage.

### Allele dropout analysis mode

We started with raw fastq files which were aligned to the GRCh37 human reference genome using BWA mem version 0.7.10 (11), the bam files were then realigned and recalibrated using GATK 3.6 (12). The clones and the single cells were compared to each other using Mutect2 (13), Strelka2 (14) and SCOUT (9). For the clones, mutations called by both Mutect2 (13) and Strelka2 (14) with depth of 10 or more reads as well as a PASS value by both callers were used as input to All^2^. For the single cell, mutation called by Mutect2 (13), Strelka2 (14) and SCOUT (9) with depth of greater than 10 reads with PASS value from all callers were used. All^2^ was run four times with four different settings as depicted in **Fig. 2**. Post All^2^, only mutation which had allele frequency of 35% or more were considered, to further filter noise introduced during clone amplification, library preparation and sequencing. The allele dropout regions for single cell were calculated using CNVpytor (15), where the entire genome was divided into 5000 base pair bins and for each bin, a likelihood score was calculated using allele frequency of SNPs within the bin. Bins were marked as allele dropout if; i) at least one SNP in the bin had allele frequency smaller than given parameter *snp_threshold* or larger than *1-snp_threshold* (in our calculations we used *snp_threshold=0*.*01*). ii) maximum likelihood allele frequency within the bin deviated from 0.5 by more than defined *threshold* parameter (we used *threshold = 0*.*1)*. iii) closest bin with heterozygous SNPs on the left side or closest bin with heterozygous SNPs on the right side was marked as dropout. Since at 50% allele frequency, the presence of the mutation is evident whether its heterozygous (when both alleles are amplified) or homozygous (where only one allele is amplified), we retain it in the score calculation despite being in a drop out region (**Fig. S4**). Single cell QC on the MDA amplified cells was performed using Scellector (8) and only one cell (cell5) was used owing to its better quality than the rest (**Fig. S5**).

### Mutation calling for lineage analyses

The files were processed the same way as the clones above. Calls were made using allele frequency cut-off of 35% to remove mutations introduced during culturing clones. Additionally, only INDELs shorter than 10bp (most confident calls) were used. Pairwise comparison between bulk data and the clones were done using consensus calls between Mutect2 and Strelka2. Mutations with a depth greater than 10 read, with at least 2 alternate supporting reads and PASS value from both callers were used. For the allele frequency plots (**Fig. 2A&C**), all mutations from All^2^, bulk, and pairwise comparison were used. For details, including calling mosaic mutation from bulk tissue and lineage tree construction please refer to the method section of Fasching et al (6).

## Availability of data

All primary data for iPSC lines we used for lineage reconstruction were accessed from at NIH NIHM Data Archives (https://nda.nih.gov/study.html?id=1057)(6). All primary data for fetal brain we used in ADA mode were accessed from NIH NIMH Data Archives(https://nda.nih.gov/edit_collection.html?id=2330) (5).

## Reference

1. Martincorena I, Fowler JC, Wabik A, Lawson ARJ, Abascal F, Hall MWJ, et al. Somatic mutant clones colonize the human esophagus with age. Science. 2018;362(6417):911–7.

2. Yokoyama A, Kakiuchi N, Yoshizato T, Nannya Y, Suzuki H, Takeuchi Y, et al. Age-related remodelling of oesophageal epithelia by mutated cancer drivers. Nature. 2019;565(7739):312–7.

3. Lee-Six H, Øbro NF, Shepherd MS, Grossmann S, Dawson K, Belmonte M, et al. Population dynamics of normal human blood inferred from somatic mutations. Nature. 2018;561(7724):473–8.

4. Zhang L, Dong X, Lee M, Maslov AY, Wang T, Vijg J. Single-cell whole-genome sequencing reveals the functional landscape of somatic mutations in B lymphocytes across the human lifespan. Proc Natl Acad Sci U S A. 2019;116(18):9014–9.

5. Bae T, Tomasini L, Mariani J, Zhou B, Roychowdhury T, Franjic D, et al. Different mutational rates and mechanisms in human cells at pregastrulation and neurogenesis. Science. 2018;359(6375):550–5.

6. Fasching L, Jang Y, Tomasi S, Schreiner J, Tomasini L, Brady MV, et al. Early developmental asymmetries in cell lineage trees in living individuals. Science. 2021;371(6535):1245–8.

7. Chen X, Schulz-Trieglaff O, Shaw R, Barnes B, Schlesinger F, Källberg M, et al. Manta: rapid detection of structural variants and indels for germline and cancer sequencing applications. Bioinformatics. 2016;32(8):1220–2.

8. Sarangi V, Jourdon A, Bae T, Panda A, Vaccarino F, Abyzov A. SCELLECTOR: ranking amplification bias in single cells using shallow sequencing. BMC Bioinformatics. 2020;21(1):521.

9. Wei J, Zhou T, Zhang X, Tian T. SCOUT: A new algorithm for the inference of pseudo-time trajectory using single-cell data. Comput Biol Chem. 2019;80:111–20.

10. Evrony GD, Cai X, Lee E, Hills LB, Elhosary PC, Lehmann HS, et al. Single-neuron sequencing analysis of L1 retrotransposition and somatic mutation in the human brain. Cell. 2012;151(3):483–96.

11. Li H, Durbin R. Fast and accurate short read alignment with Burrows-Wheeler transform. Bioinformatics. 2009;25(14):1754–60.

12. Poplin R, Ruano-Rubio V, DePristo MA, Fennell TJ, Carneiro MO, Van der Auwera GA, et al. Scaling accurate genetic variant discovery to tens of thousands of samples. bioRxiv. 2018:201178.

13. Cibulskis K, Lawrence MS, Carter SL, Sivachenko A, Jaffe D, Sougnez C, et al. Sensitive detection of somatic point mutations in impure and heterogeneous cancer samples. Nat Biotechnol. 2013;31(3):213–9.

14. Saunders CT, Wong WS, Swamy S, Becq J, Murray LJ, Cheetham RK. Strelka: accurate somatic small-variant calling from sequenced tumor-normal sample pairs. Bioinformatics. 2012;28(14):1811–7.

15. Suvakov M, Panda A, Diesh C, Holmes I, Abyzov A. CNVpytor: a tool for CNV/CNA detection and analysis from read depth and allele imbalance in whole genome sequencing. bioRxiv. 2021:2021.01.27.428472.

